# FlyORF-TaDa allows rapid generation of new lines for *in vivo* cell-type specific profiling of protein-DNA interactions in *Drosophila melanogaster*

**DOI:** 10.1101/2020.08.06.239251

**Authors:** Gabriel N Aughey, Caroline Delandre, Tony D Southall, Owen J Marshall

## Abstract

Targeted DamID (TaDa) is an increasingly popular method of generating cell-type specific DNA binding profiles *in vivo*. Although sensitive and versatile, TaDa requires the generation of new transgenic fly lines for every protein that is profiled, which is both time-consuming and costly. Here, we describe the FlyORF-TaDa system for converting an existing FlyORF library of inducible open reading frames (ORFs) to TaDa lines via a genetic cross, with recombinant progeny easily identifiable by eye colour. Profiling the binding of the H3K36me3-associated chromatin protein MRG15 in larval neural stem cells using both FlyORF-TaDa and conventional TaDa demonstrates that new lines generated using this system provide accurate and highly-reproducible DamID binding profiles. Our data further show that MRG15 binds to a subset of active chromatin domains *in vivo*. Courtesy of the large coverage of the FlyORF library, the FlyORF-TaDa system enables the easy creation of TaDa lines for 74% of all transcription factors and chromatin modifying proteins within the *Drosophila* genome.

## 1 Introduction

Characterising the specific protein-DNA interactions that underlie gene expression is essential for understanding the biology of any given tissue. The last decade has seen a change in thinking regarding the action of transcription factors (TFs), proteins that bind to enhancer and promoter regions of genes and modify the level of gene expression. In particular, it is now established that TFs work in complex groups or communities to control gene expression (for review, see Reiter et al., 2017). A key recent finding has been that the majority of TFs can act as either an activator or as a repressor, with their function determined by the surrounding TF community (Stampfel et al., 2015). As such, the ability to profile modules of TFs is critical to understanding regulatory function.

A major challenge in undertaking the systematic profiling of TF binding within a cell is the availability of reagents. For ChIP-seq, a lack of appropriate antibodies is a considerable impediment, combined with the difficulty of profiling TFs that are not directly bound to DNA. These issues are solved by DamID, which effectively profiles all TFs coming into proximity with DNA and requires no antibodies for profiling (for review, see (Aughey et al., 2019)).

Targeted DamID (TaDa) allows DamID to be applied in a cell-type specific manner to profile cellular transcriptional machinery (Southall et al., 2013), chromatin modifying proteins (Marshall and Brand, 2017), nuclear structural proteins and transcription factors (Doupé et al., 2018). The technique allows the binding profile of any protein associated with DNA to be mapped in living organisms without the concern of fixation-induced artefacts, from very small amounts of material (10,000 cells being enough to generate high-quality profiles) and without the need for cell-sorting (Marshall et al., 2016). Profiling is performed *in vivo* using the GAL4/UAS system (Brand and Perrimon, 1993) to provide cell-type specificity. As the GAL4/UAS system drives strong expression of transgenes, and high levels of Dam are toxic in eukaryotic cells, TaDa uses a bicistronic transcript to greatly reduce the translation of Dam-fusion proteins. In this system, a long primary ORF is separated from the Dam-fusion ORF by two stop codons and a frame shift, with translation of the latter arising through low rates of spontaneous ribosomal re-initiation. To generate DNA-binding profiles, GAL4 driver lines are crossed to lines carrying TaDa Dam-fusion proteins. However, the time and costs of generating the transgenic fly lines required for TaDa profiling are considerable.

The FlyORF library of fly lines (Bischof et al., 2013) contains inducible open reading frames (ORFs) that cover 74% (560/757) of all known and predicted TFs and chromatin-associated proteins in the *Drosophila* genome (Bischof et al., 2013, 2014). The library was designed with cassette exchange features, presenting the possibility of converting existing FlyORF lines into TaDa lines to profile DNA binding via a simple cross. Here, we describe FlyORF-TaDa, a new system that allows conversion of FlyORF lines to TaDa lines via Flippase-mediated cassette exchange. FlyORF-TaDa permits the rapid and easy generation of new TF-binding profiles without cloning or the creation of transgenic animals, with recombinants easily identified by eye colour and fluorescence.

## 2 Materials and Methods

### 2.1 Fly husbandry

*Drosophila* were raised on media containing 5% (w/v) yeast, 5.5% (w/v) dextrose, 3.5% (w/v) cornflour, 0.75% (w/v) agar, 0.25% (v/v) Nipagin and 0.4% (v/v) propionic acid. Flies were grown in incubators at 70% humidity on a 12 hour / 12 hour light / dark cycle.

### 2.2 Expression constructs

The *pFlyORF-TaDa* vector was created by cutting *pTaDaG2* (Delandre et al., 2020) with BglII/NotI (NEB) and inserting a synthetic gBlock (IDT) containing an *FRT5* site followed by an in-frame stop codon via NEB HiFi Assembly (NEB) to create *pTaDaG2-FRT5*. *pTaDaG2-FRT5* was cut with AfeI/BstBI and a synthetic gBlock (IDT) containing *3xP3-DsRed2-small_t_intron-polyA* via NEB Hifi Assembly to generate *pTaDaG2-FRT5-3xP3-dsRed2*. Finally, mini-white (including the 240 bp upstream and 630 bp downstream *white* regulatory regions) was amplified via PCR from *pTaDaG2* and inserted into *pTaDaG2-FRT5-3xP3-dsRed* cut with XhoI and XbaI via NEB Hifi Assembly to generate *pFlyORF-TaDa*.

*pTaDaG2-MRG15* was generated by inserting a synthetic gBlock (IDT) containing the sequence of MRG15-RA into the *pTaDaG2* vector cut with XbaI/XhoI.

All plasmids were sequence-verified via Sanger sequencing (ABI).

Plasmid maps were generated using SnapGene software (Insightful Science).

### 2.3 Fly lines

The *worniu-GAL4*; *tub-GAL80ts* line (Albertson et al., 2004) was used as a driver for neural stem cells. The *FlyORF-TaDa* line was created through phiC31-integrase-mediated insertion into *ZH-86FB* (the site used by the FlyORF expression library) on chromosome 3, by injecting *pFlyORF-TaDa* at 200ng/µL into *y[1] M{RFP[3xP3.PB] GFP[E.3xP3]=vas-int.Dm}ZH-2A w[*]; M{3xP3-RFP.attP}ZH-86Fb* (Bloomington #24749) embyros. The resulting transgenic line was backcrossed three times to *w*^*1118*^ before a homozygous stock was generated. The *w; hs-FlPD5; FlyORF-TaDa* was generated by crossing to *w*^*1118*^; *P{y[+t7.7] w[+mC]=hs-FLPD5}attP40/CyO* stock (Bloomington #55812). The *TaDaG2-MRG15* line was generated by BestGene, Inc (CA), through phiC31-integrase-mediated insertion of *pTaDaG2-MRG15* into *attP2* on chromosome 3L. *TaDaG2-Dam* flies were used as previously published (Delandre et al., 2020).

### 2.4 FlyORF cassette exchange

Homozygous *hs-FlpD5; FlyORF-TaDa* virgin females were crossed to males from *FlyORF* lines. Progeny (third instar, 96 hours after larval hatching) were heat shocked at 37℃ for 45 mins. After eclosion, F1 male flies were crossed to TM3/TM6B virgin females. F2 males and virgin females with TM6B and exhibiting the correct eye phenotype (*w-; dsRed+*) were crossed to establish a balanced stock.

### 2.5 Targeted DamID

Appropriate lines (for conventional TaDa: *TaDaG2-MRG15* and *TaDaG2-Dam*; for FlyORF-TaDa: *FlyORF-TaDa-MRG15* and *FlyORF-TaDa* lines) were crossed to *worniu-GAL4; tub-GAL80ts* virgin females in cages. Embryos were collected on apple juice agar plates with yeast over a 4 hour collection window and grown at 18°C for two days. Newly hatched larvae were transferred to food plates for a further five days at 18°C, before shifting to 29°C for 24 hours.

Larval brains were dissected in PBS, and processed for DamID-seq as previously described (Marshall et al., 2016; Marshall and Brand, 2017) with the following modifications. Briefly, DNA was extracted using a Quick-DNA Miniprep plus kit (Zymo), digested with DpnI (NEB) overnight and cleaned-up with a PCR purification kit (Machery-Nagel), DamID adaptors were ligated, digested with DpnII (NEB) for 2 hours, and amplified via PCR using MyTaq DNA polymerase (Bioline).

Following amplification, 2 µg DNA was sonicated in a Bioruptor Plus (Diagenode). DamID adaptors were removed by AlwI digestion, and 500 ng of the resulting fragments end-repaired with a mix of enzymes (T4 DNA ligase (NEB) + Klenow Fragment (NEB) + T4 polynucleotide kinase (NEB)), A-tailed with Klenow 3’ to 5’ exo-(NEB), ligated to Illumina Truseq LT adaptors using Quick Ligase enzyme (NEB) and amplified via PCR with NEBNext Hi-fidelity enzyme (NEB). The resulting next-generation sequencing libraries were sequenced on a HiSeq2500 (Illumina).

### 2.6 Bioinformatic analyses

DamID binding profiles were generated from NGS reads using damidseq_pipeline (Marshall and Brand, 2015) and visualised using pyGenomeTracks (Ramírez et al., 2018). Peaks were called using a three-state hidden markov model via the hmm.peak.caller R script (freely available from https://github.com/owenjm/hmm.peak.caller).

Heatmaps were generated via the ComplexHeatmap R package (Gu et al., 2016). Gene Ontology (GO) enrichment plots were performed using the ClusterProfiler R package with Bonferroni-Holm adjusted *P* values; enrichmap plots were generated by limiting GO terms to <1000 genes and using the simply() function to remove redundancy (Yu et al., 2012).

MRG15 enrichment by chromatin state (significance and odds ratio) was assessed via Fisher’s exact test with an alternate hypothesis of “greater”, for contingency tables of genomic coverage of MRG15 vs genomic coverage of the chromatin state in NSCs. All *P* values were Bonferroni-Holm adjusted.

All other plots were generated using R (R Development Core Team, 2011).

### 2.7 Data availability

The sequence of *pFlyORF-TaDa* is available under Genbank accession number MT733231. DamID-seq data and processed bedgraph files are available upon request and will be deposited in NCBI GEO.

## 3 Results and Discussion

### 3.1 Design of the FlyORF-TaDa system

The FlyORF library of inducible open reading frames (ORFs) incorporates an FRT5 site immediately upstream of each ORF, allowing Flippase-mediated exchange with an upstream donor cassette in the same genomic insertion site (Bischof et al., 2013). To convert these lines to TaDa lines, FlyORF-TaDa provides the UAS-inducible bicistronic transcript of TaDa vectors upstream of an FRT5 site as the donor cassette. FlyORF-TaDa uses the TaDaG version of the TaDa bicistronic transcript, in which the primary ORF is myristoylated GFP, providing the advantage of easily-detectable fluorescent labelling of cells being profiled within experimental samples (Delandre et al., 2020).

The bicistronic cassette is followed by an *FRT5* site in frame with *Dam* (Fig. 1A; Fig. S1A). Without recombination, a stop codon positioned directly after the FRT5 site allows the FlyORF-TaDa line to be used as a Dam-only control for DamID signal normalization (Fig. S1B). Following recombination with a FlyORF line, the FRT5 site together with the Gateway cloning residual attB1 site present as part of the FlyORF library construction (Bischof et al., 2013) becomes a 23 amino acid protein linker region (Fig. S1C).

**Figure 1.**
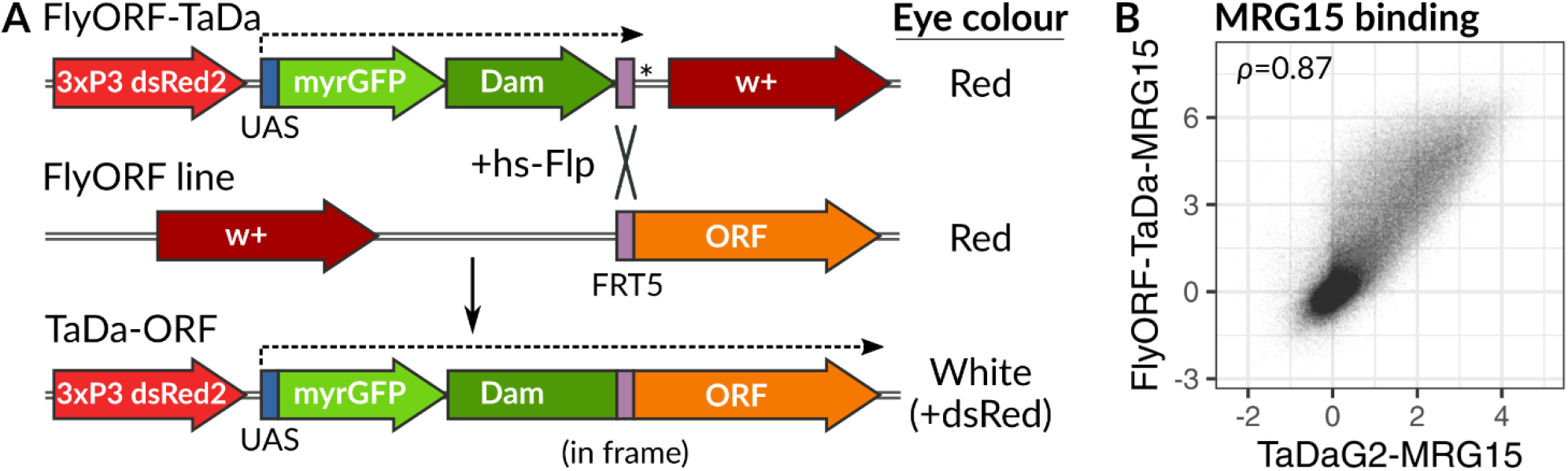
The FlyORF-TaDa system generates new lines for cell-type specific profiling. (A) Schematic of the FlyORF-TaDa system. When the FlyORF-TaDa donor line is crossed to a FlyORF line and a heat-shock-inducible *flippase* (hs-Flp), recombination between compatible FRT5 sites generates a new TaDa line for the ORF in the progeny. These lines can be screened for in the subsequent generation by eye colour. Dashed lines show GAL4/UAS-induced transcription; * = stop codon. (B) Correlation of DNA binding profiles generated for MRG15 in NSCs using the FlyORF-TaDa system, compared to conventional TaDa (Pearson’s correlation is shown).

To prevent the need to PCR-screen progeny, the vector was additionally designed with a *3xP3-dsRed2* eye marker upstream of the TaDa cassette, and the *mini-white* eye marker downstream of the *FRT5* site (Fig. 1A, Fig. S1A). While both parental strains used for a conversion are *w+*, upon Flippase-mediated strand exchange, successful recombinants will be *dsRed2+, w-* making screening of recombinants by eye colour fast and straightforward.

The *pFlyORF-TaDa* vector was inserted into ZH-86FB, the same insertion site used by the FlyORF library, and the resulting line was crossed with a heat-shock-inducible Flippase (*hs-FlpD5*) line to yield an *hs-FlpD5; FlyORF-TaDa* donor line. TaDa alleles are generated by crossing any FlyORF line to this donor line, in conjunction with a heat-shock of progeny during the larval phase. An illustration of the crossing scheme is shown in Fig. S2.

### 3.2 Profiling of MRG15 binding in Neural Stem Cells using the FlyORF-TaDa system

To determine whether FlyORF-TaDa lines generated through the system faithfully profiled binding in a cell-type specific manner, we obtained binding profiles for the chromatin binding protein MRG15 in neural stem cells (NSCs). We obtained profiles from a *FlyORF-TaDa-MRG15* and compared these to MRG15 profiles generated using conventional TaDa, driving expression in both cases with the NSC-specific driver, *worniu-GAL4*. Excellent correlation between samples was observed throughout (Fig. 1B,C), both between individual replicates (Fig. S3) and when comparing the average profiles from the two systems (Fig. 1B,C). In particular, the two biological replicates generated using the FlyORF-TaDa system showed extremely high reproducibility (Pearson’s correlation between replicates, 0.95; Fig. S3).

MRG15 is a protein associated with the H3K36me3 histone mark (Zhang et al., 2006), which in turn is associated with transcribed exons (Kolasinska-Zwierz et al., 2009; Kharchenko et al., 2011). The protein is also a key component of the “Yellow” active chromatin state described in flies by (Filion et al., 2010), a study which reduced chromatin configurations in *Drosophila* to five broad classification types, or colours. In concordance with these observations, we found MRG15 bound at previously-published Yellow chromatin domains in NSCs (Marshall and Brand, 2017) (Fig. 2A), with MRG15-bound peaks covering 80% (9.8Mb of 12.2Mb) of Yellow chromatin. Whilst MRG15 occupancy was significantly enriched to some degree over all three active chromatin states in NSCs, by far the greatest enrichment was observed over Yellow chromatin (Fig. 2B; log_*e*_θ (Yellow chromatin) = 2.91, Fisher’s exact test).

**Figure 2.**
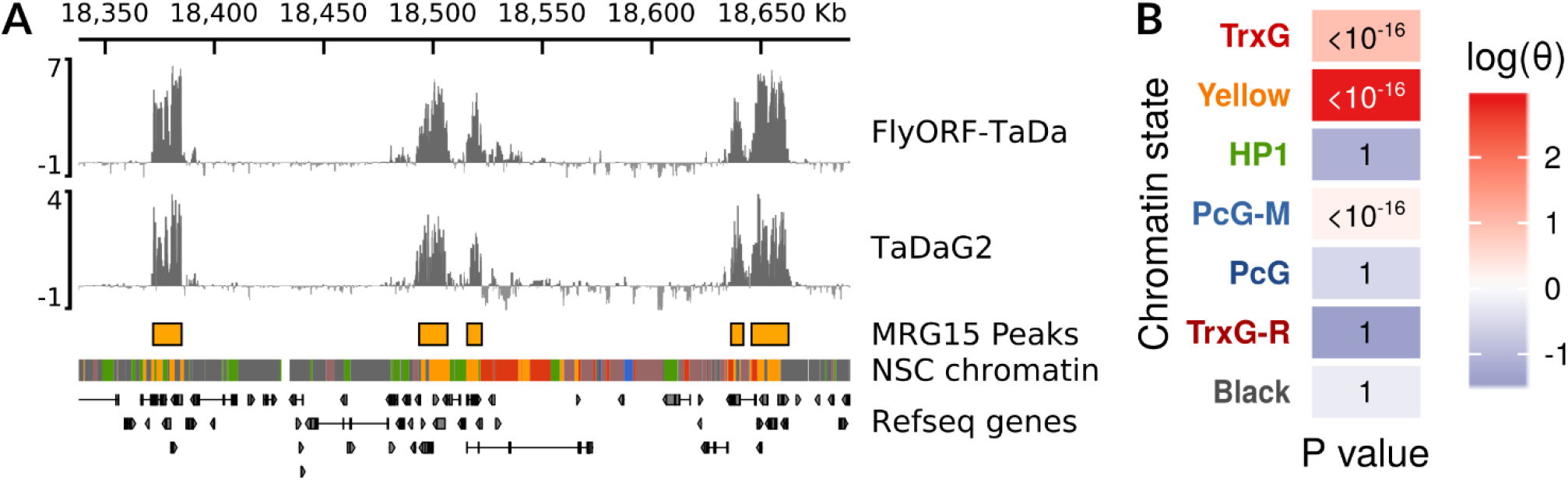
Characteristics of MRG15 binding in NSCs. (A) MRG15 binds to regions of Yellow chromatin in NSCs. Profiles obtained via FlyORF-TaDa and conventional TaDa are shown; peaks were called on the average binding profile of the combined MRG15 datasets. NSC chromatin state data and colours are from Marshall and Brand (2017). A 300Kb region of chrX is illustrated. (B) Significance and odds ratio (θ) of enrichment of MRG15 peak occupancy within NSC chromatin states as assessed via Fisher’s exact test.

Genes covered by bound peaks were enriched for Gene Ontology functions associated with metabolism (Fig. 3, Fig. S4), a known association for genes covered by Yellow chromatin, both in cell lines (Filion et al., 2010; Bemmel et al., 2013) and in NSCs (Marshall and Brand, 2017). Importantly, we also observed an enrichment for genes involved in nervous system development expressed in NSCs (Fig. 3), consistent with occupancy over transcribed exons and indicating that the FlyORF-TaDa system can faithfully profile cell-type specific protein binding *in vivo*.

**Figure 3.**
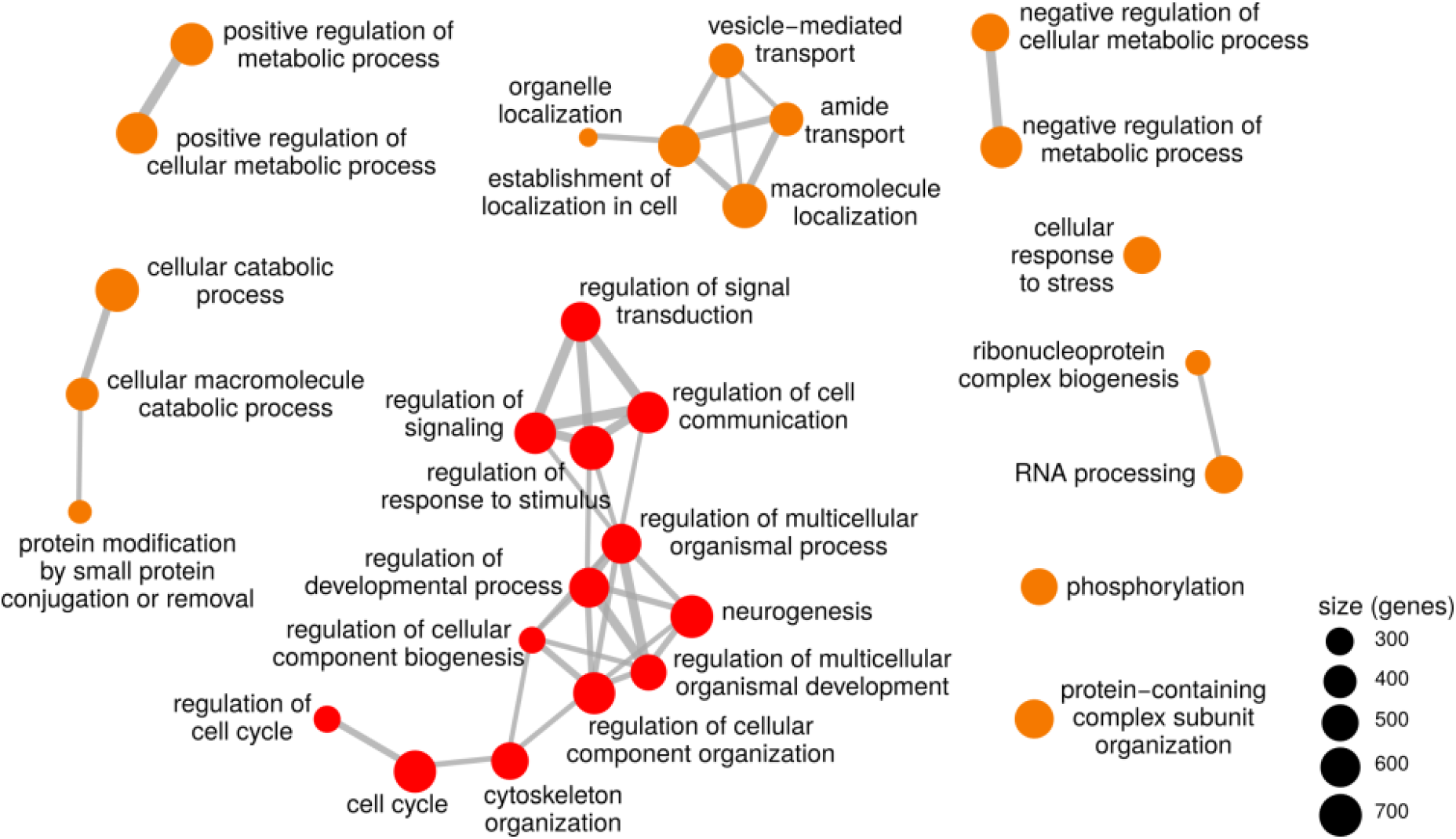
Gene Ontology (GO) term enrichment for genes covering MRG15 bound peaks identifies terms for both metabolism and developmental neurogenesis (non-redundant GO terms covering <1000 genes illustrated). Metabolism-related terms are coloured in orange; developmental terms are coloured in red. All terms illustrated were significantly enriched (all adjusted *P* values < 10^-28^).

### 3.3 Conclusion

The FlyORF-TaDa system places fast and straightforward cell-type specific profiling of TF binding within the reach of any fly lab, allowing the profiling of over 74% of all TFs and chromatin-associated proteins via a simple genetic cross. Establishment of new recombinant lines from the parental FlyORF stocks is achieved in two generations, without the need for cloning, sequencing or transgenesis.

With three-quarters of all TFs covered by donor FlyORF lines, comprehensive wide-scale cell-type specific profiling of TF-binding networks can be achieved. TaDa is ideally suited to profiling such networks without introducing the variability of different fixation and antibody isolation steps found in alternative methods such as ChIP-seq. The FlyORF-TaDa system now eliminates a major obstacle to this approach by dramatically reducing the time and costs required to generate the lines required to profile TF networks. To further this aim, we have successfully generated 17 new FlyORF-TaDa lines using the system and are creating a library of converted TF and chromatin factor lines that will be deposited in stages at the Bloomington Drosophila Stock Center. We anticipate that this resource will prove highly useful to the *Drosophila* transcription and chromatin, and developmental biology communities.

## 4 Acknowledgments

We thank G. Jefferies for technical support. This work was supported by NHMRC grants APP1128784 and APP1185220 to O.J.M., a Wellcome Trust Investigator grant 104567 to T.D.S and a BBSRC grant BB/P017924/1 to T.D.S. and G.N.A.

## Supplementary figures

**Figure S1.**
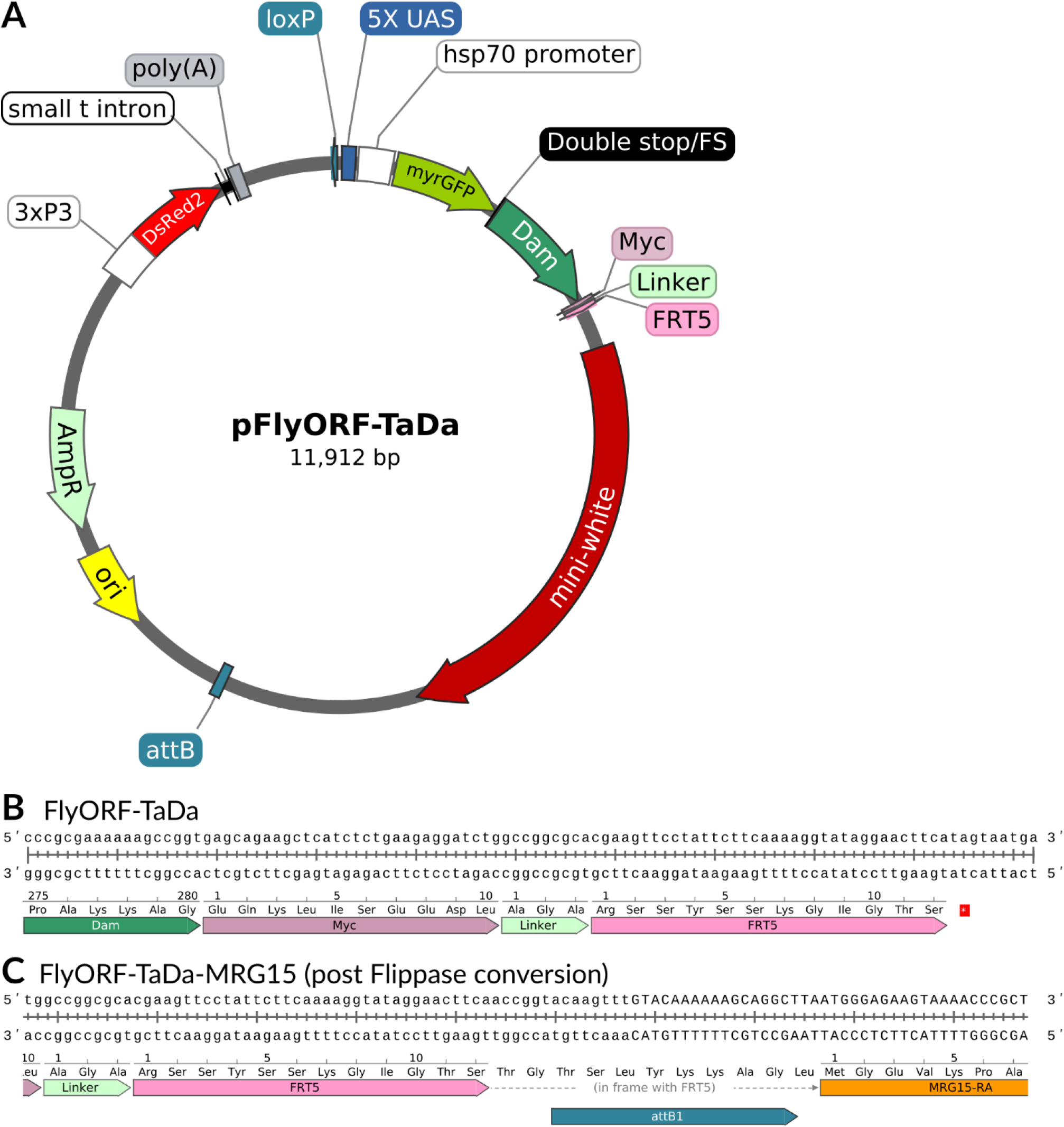
Maps of the *pFlyORF-TaDa* plasmid and converted genomic sequence. (A) Overall plasmid map schematic, with all relevant elements illustrated. (B) DNA sequence and in-frame translation of the linker region following *Dam* in the *FlyORF-TaDa* line; the STOP codon after the FRT5 site is marked with an asterisk. (C) Genomic sequence following Flippase-mediated conversion of the *FlyORF-TaDa* donor line to *FlyORF-TaDa-MRG15*.

**Figure S2.**
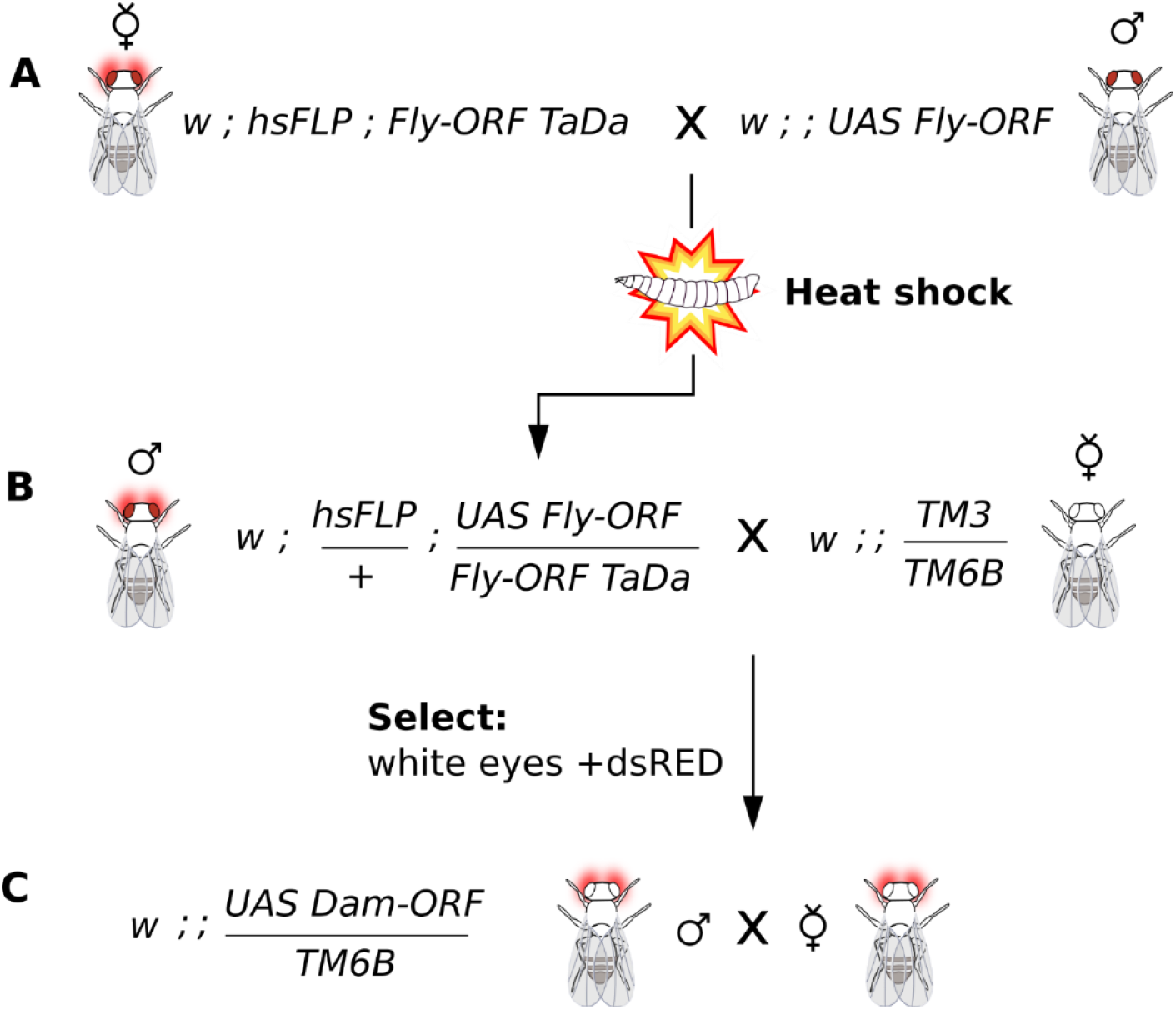
Suggested crossing scheme for generating Dam-fusion lines using FlyORF-TaDa. A) Homozygous FlyORF flies are crossed to Fly-ORF TaDa line containing hsFLP. B) F1 progeny are heat shocked during larval stages and adults crossed to third chromosome balancer. NB: Any w-line can be substituted at this stage if it is not necessary to balance immediately (e.g. if isogenising is intended). C) Resulting progeny with the correct marker combination (white eye + dsRED) over the same balancer chromosome are crossed to obtain balanced progeny for the generation of a stable stock. NB: it is expected that resulting Fly-ORF TaDa fusions will be viable as homozygotes, so will not require the presence of the balancer for long-term viability.

**Figure S3.**
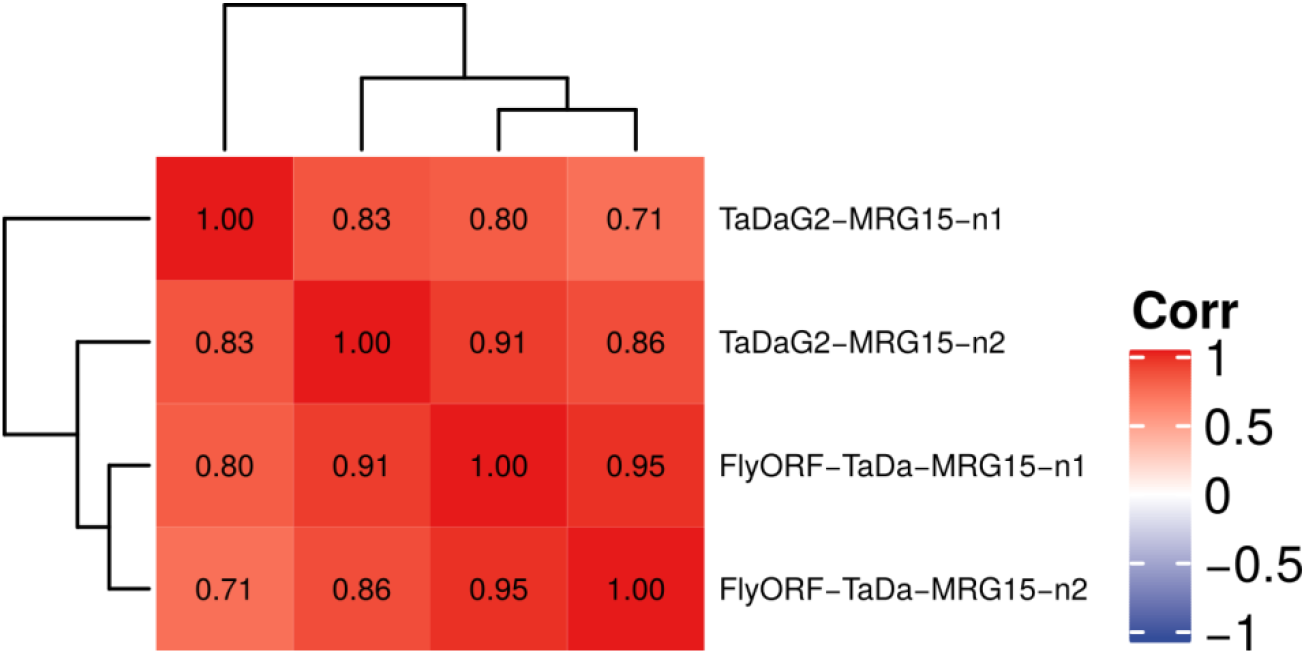
Correlation between individual replicates of MRG15 binding in NSCs acquired using either the conventional TaDa (TaDaG2) or FlyORF-TaDa. Pearson’s correlation is shown.

**Figure S4.**
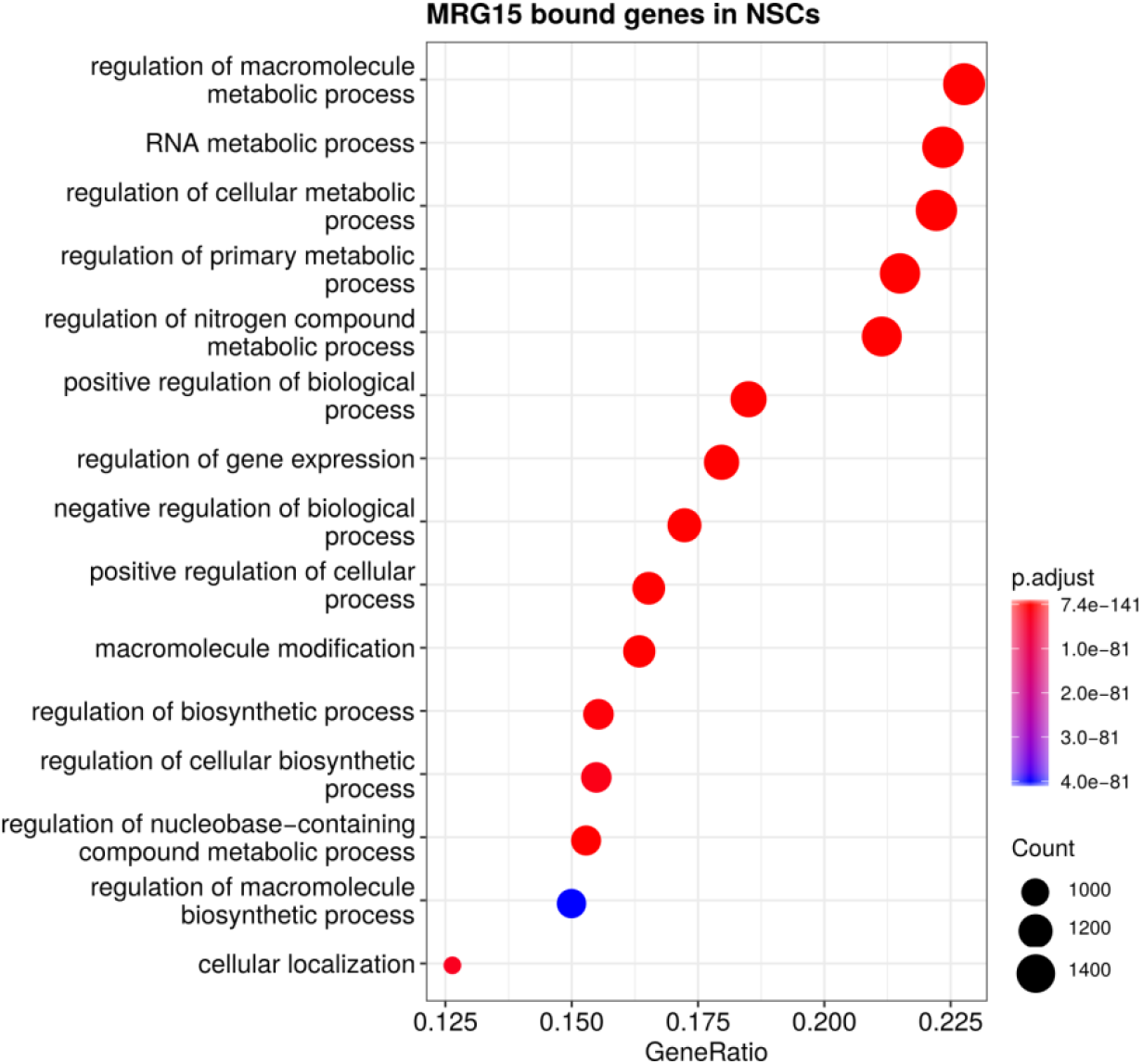
Gene Ontology (GO) term enrichment for genes covering MRG15 bound peaks in NSCs, with terms limited to 2000 genes.

